# An infectivity-enhancing site on the SARS-CoV-2 spike protein is targeted by COVID-19 patient antibodies

**DOI:** 10.1101/2020.12.18.423358

**Authors:** Yafei Liu, Wai Tuck Soh, Asa Tada, Akemi Arakawa, Sumiko Matsuoka, Emi E. Nakayama, Songling Li, Chikako Ono, Shiho Torii, Kazuki Kishida, Hui Jin, Wataru Nakai, Noriko Arase, Atsushi Nakagawa, Yasuhiro Shindo, Masako Kohyama, Hironori Nakagami, Keisuke Tomii, Koichiro Ohmura, Shiro Ohshima, Masato Okada, Yoshiharu Matsuura, Daron M. Standley, Tatsuo Shioda, Hisashi Arase

## Abstract

SARS-CoV-2 infection causes severe symptoms in a subset of patients, suggesting the presence of certain unknown risk factors. Although antibodies against the receptor-binding domain (RBD) of the SARS-CoV-2 spike have been shown prevent SARS-CoV-2 infection, the effects of antibodies against other spike protein domains are largely unknown. Here, we screened a series of anti-spike monoclonal antibodies from COVID-19 patients, and found that some of antibodies against the N-terminal domain (NTD) dramatically enhanced the binding capacity of the spike protein to ACE2, and thus increased SARS-CoV2 infectivity. Surprisingly, mutational analysis revealed that all the infectivity-enhancing antibodies recognized a specific site on the surface of the NTD. The antibodies against this infectivity-enhancing site were detected in all samples of hospitalized COVID-19 patients in the study. However, the ratio of infectivity-enhancing antibodies to neutralizing antibodies differed among patients. Furthermore, the antibodies against the infectivity-enhancing site were detected in 3 out of 48 uninfected donors, albeit at low levels. These findings suggest that the production of antibodies against SARS-CoV-2 infectivity-enhancing site could be considered as a possible exacerbating factors for COVID-19 and that a spike protein lacking such antibody epitopes may be required for safe vaccine development, especially for individuals with pre-existing enhancing antibodies.

## Main

SARS-CoV-2 is a novel coronavirus that causes coronavirus disease 2019 (COVID-19)^1^. Although SARS-CoV-2 infection can result in severe symptoms and is associated with high mortality in some patients, most infected individuals do not exhibit such severe symptoms; therefore, additional factors are likely to facilitate the progression to severe COVID-19^2,3^. SARS-CoV-2, an enveloped positive-strand RNA virus, requires fusion with the host cell membrane for infection^4^. The spike protein is the major envelope protein in SARS-CoV-2 and is composed of S1 and S2 subunits. The S1 subunit is further divided into an N-terminal domain (NTD) and a receptor-binding domain (RBD)^5,6^. The interaction between the RBD and the host cell receptor, ACE2, is responsible for SARS-CoV-2 infection of host cells^7,8^.

COVID-19 patients produce antibodies against the RBD of the spike protein, blocking the SARS-CoV-2 infection^9-11^. Therefore, antibody production against the spike protein plays a pivotal role in host defense against SARS-CoV-2 infection^9-11^. However, antibodies against viruses are not always protective^12^. For example, antibodies against dengue virus protein can cause severe diseases that are mediated by the Fc receptor^13^. In this study, we evaluated the effect of various types of anti-spike antibodies on ACE2 binding and SARS-CoV-2 infection.

## Results

### Enhanced ACE2 binding to the spike protein by a subset of anti-NTD antibodies

We studied the function of the antibodies produced in COVID-19 patients by generating a series of anti-spike monoclonal antibodies from COVID-19 patients^9-11,14^ and analyzing their effect on the binding of recombinant ACE2 to cells expressing the spike protein (Fig. 1a). The binding specificity of the antibodies was determined using transfectants encoding the membrane form of the NTD, RBD, or S2 domain of the spike protein (Supplementary Fig. 1a,b). Some anti-spike antibodies that did not bind to recombinant NTD in previous reports^11,14^ specifically recognized NTD expressed on the cell surface. Recombinant ACE2 bound to whole spike and RBD-TM transfectants but not to NTD-TM or mock transfectants, indicating that recombinant ACE2 specifically binds to the RBD of the spike protein (Supplementary Fig. 2). As expected, most of the antibodies against the RBD domain blocked the binding of ACE2 to the spike protein, and antibodies against the S2 subunit did not affect ACE2 binding. However, a specific subset of antibodies—8D2^14^, 2210^11^, 2369^11^, 2490^11^, 2582^11^, and 2660^11^—targeting the NTD domain (denoted “enhancing antibodies”) were found to enhance the binding of ACE2 to the spike protein (Fig. 1a and Supplementary Fig. 2). By contrast, other anti-NTD antibodies, such as 2016^11^ and 4A8^14^, did not affect the binding of ACE2 to the spike protein, although they recognized the full-length spike protein and the NTD domain to a similar degreee as the enhancing antibodies (Fig. 1a and Supplementary Fig. 1b). The increased ACE2 binding by the enhancing antibodies the was not observed in RBD-TM or NTD-TM transfectants (Supplementary Fig. 2). Furthermore, the enhancement of ACE2 binding to the spike protein by the enhancing antibodies was dose-dependent (Fig. 1b).

**Fig. 1.**
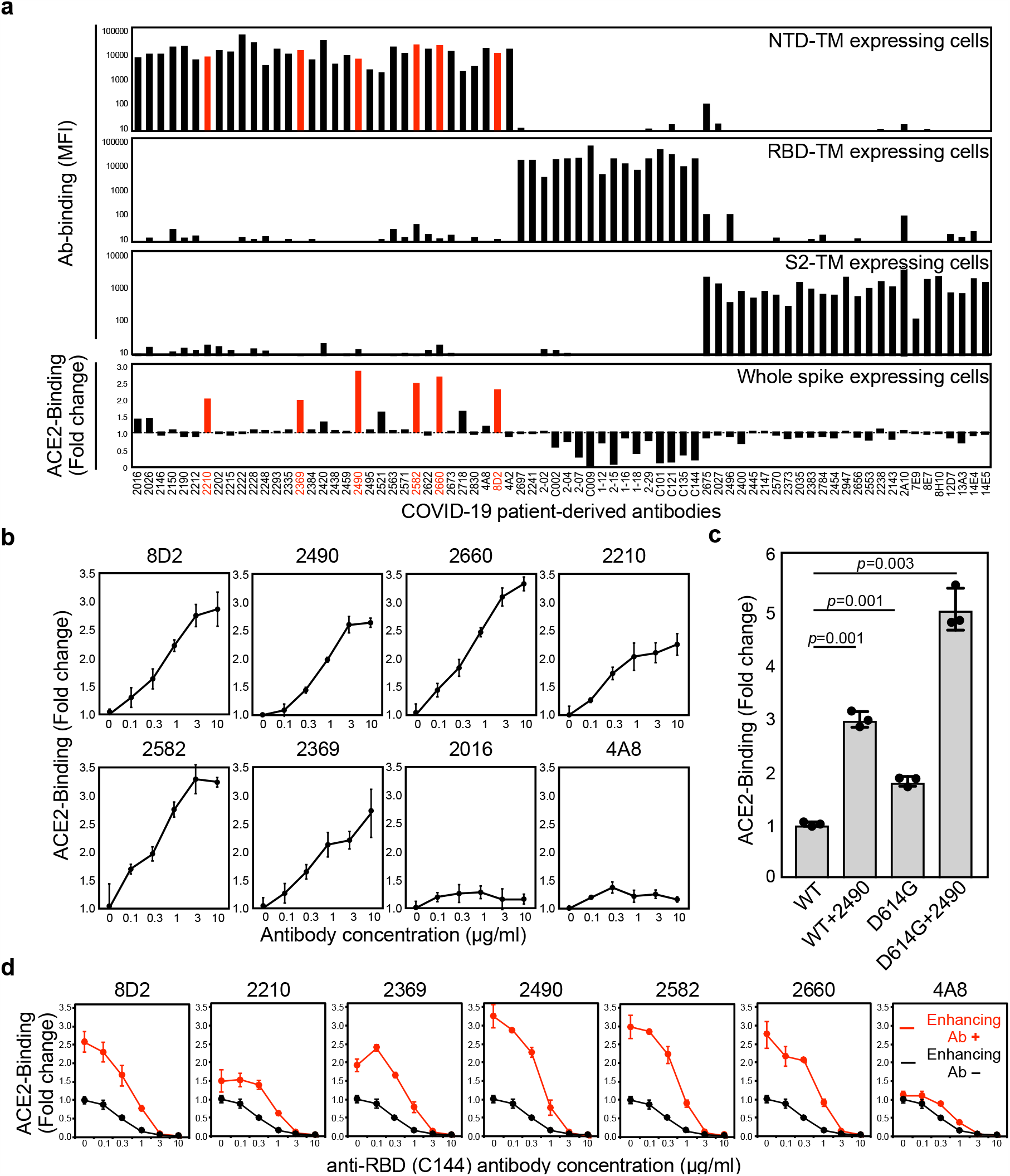
Enhanced ACE2 binding to spike protein by some of anti-NTD antibodies. **a**, The HEK293 cells transfected with vectors expressing NTD-TM, RBD-TM, and S2-TM were stained with anti-spike antibodies. The mean fluorescence intensity (MFI) of the stained cells was calculated (top three columns). The binding of ACE2-Fc-fusion protein to full-length spike transfectants was analyzed in the presence of the indicated antibodies at 1 μg/ml (bottom column). The antibodies that enhanced ACE2-Fc binding to the spike transfectants by more than 1.9 times are indicated in red. **b**, ACE2-Fc binding to the spike transfectants in the various concentrations of antibodies. **c**, ACE2-Fc binding to the wild-type or D614G spike protein in the presence of 3 μg/ml of 2490 mAb. The statistical significance derived from an unpaired *t*-test is indicated. **d**, ACE2-Fc binding to wild-type spike protein in the presence of the indicated antibodies at 3 μg/ml and various concentrations of anti-RBD neutralizing antibody C144 (red line). ACE2-Fc binding in the absence of the enhancing antibodies was shown as the control (black line). The data from triplicates are presented as mean ± SD. The representative data from three independent experiments are shown.

The D614G mutant of the spike protein, discovered in recently isolated strains of SARS-CoV-2, is rapidly spreading because of its higher infectivity than that of the wild-type spike protein^15-20^. Recent structural analysis indicated that ACE2 binding domain of the RBD is more exposed in the D614G mutant spike protein than in the wild-type spike protein^15-17^. Compatible with this observation, recombinant ACE2 bound to the D614G spike protein more than to the wild-type protein, although the cell surface expression of the spike proteins were comparable (Fig 1c and Supplementary Fig. 1c). Notably, the effect of the enhancing antibodies on ACE2 binding to the spike protein was higher than that of the D614G mutation^21^. Moreover, the enhancing antibodies increased the binding of ACE2 to the D614G spike protein.

Since anti-RBD antibodies that block the binding of ACE2 to the spike protein play a central role in antibody-mediated host defense against SARS-CoV-2 infection^9-11^, we analyzed how the enhancing antibodies affected the ACE2-blocking activity of a representative anti-RBD antibody C144^10^ (Fig. 1d). Interestingly, the binding of ACE2 to the spike protein increased in the presence of the enhancing antibodies, even upon addition of C144 at concentrations up to 1 μg/ml, suggesting that the neutralization activity of the anti-RBD antibodies is reduced in the presence of the enhancing antibodies.

### The enhancing antibodies facilitate SARS-CoV-2 infectivity

The effect of enhancing antibodies on ACE2 binding to the spike protein suggested that the infectivity of SARS-CoV-2 would also be increased, similar to the effect of the D614G mutation^16,17,19,20^. We utilized vesicular stomatitis virus (VSV)/ΔG-GFP SARS-CoV-2 spike pseudovirus (SARS-CoV-2 PV) to analyze the effect of representative enhancing antibodies on SARS-CoV-2 infection. The enhancing antibodies increased the infectivity of SARS-CoV-2 PV to the ACE2-transfected HEK293 cells in a enhancing antibody dose-dependent manner (Fig. 2a and Supplementary Fig. 3). In contrast, an anti-NTD antibody (4A2) that did not enhance ACE2 binding to the spike protein (Fig. 1b,d) did not increase the infectivity. The enhanced infectivity by the antibodies was observed regardless of the amount of the virus (Fig. 2b).

**Fig. 2.**
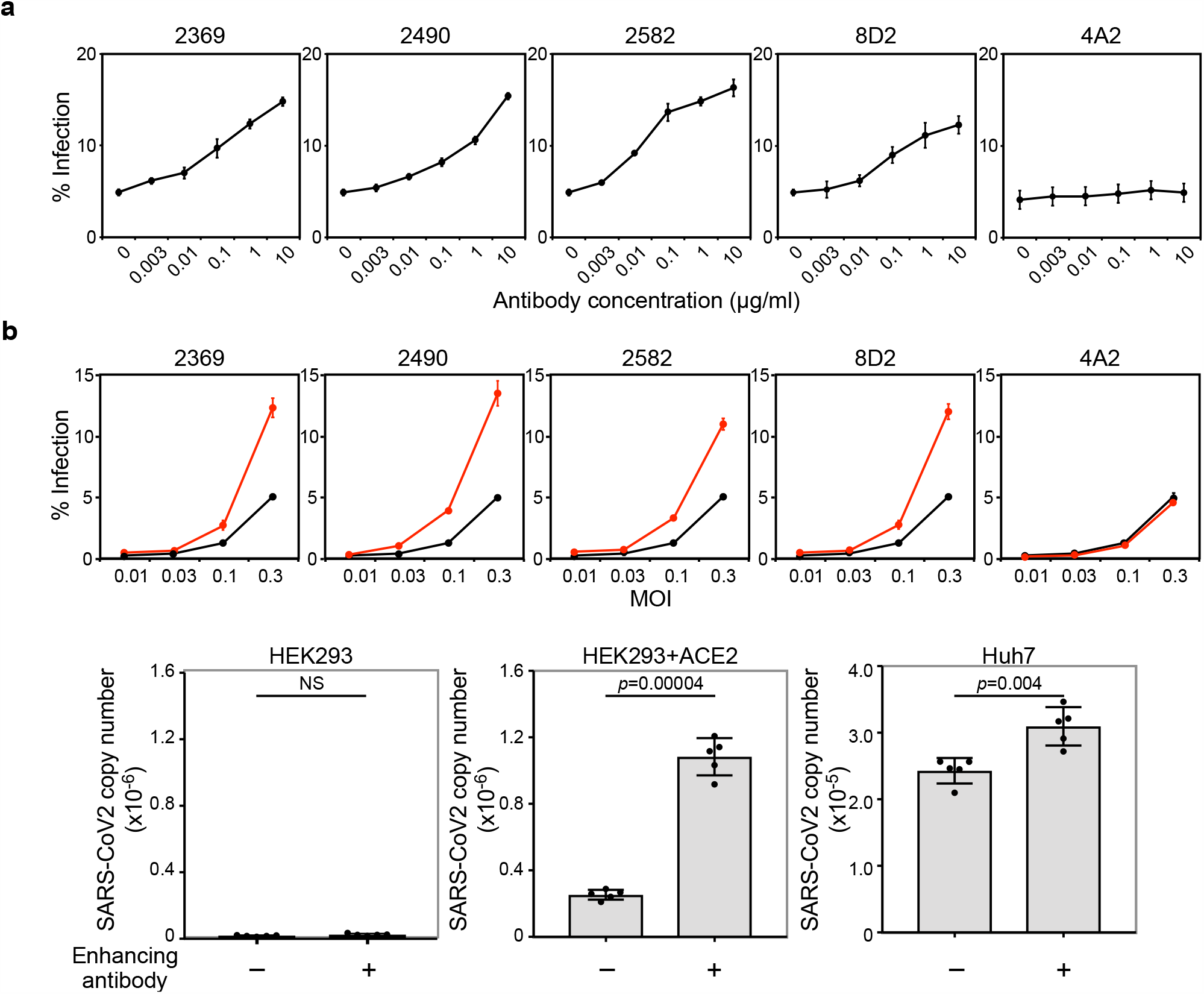
Enhanced SARS-CoV-2 infectivity by specific anti-NTD antibodies. **a**, The *ACE2*-expressing HEK293 cells (MOI: 0.3) were infected by a SARS-CoV-2-spike pseudovirus carrying a GFP reporter gene in the presence of various concentrations of indicated antibodies. The proportion of GFP-positive infected cells is shown. **b**, The *ACE2*-expressing HEK293 cells were infected with a SARS-CoV-2-spike pseudovirus carrying a GFP reporter gene at different MOIs with (red line, 3 μg/ml) or without (black line) the indicated antibodies. **c**, HEK293 cells, ACE2-expressing HEK293 cells, and Huh7 cells were infected with authentic SARS-CoV-2 virus in the presence (+) or absence (–) of enhancing antibody 2490 at 1 μg/ml. The amounts of SARS-CoV-2 virus produced in the cell culture supernatants were analyzed 48 h after infection. The statistical significance derived from an unpaired *t*-test is indicated. NS: Not significant. The data are presented as mean ± SD. The representative data from three independent experiments are shown.

Next, we examined the effect of the enhancing antibodies using authentic SARS-CoV-2 virus. The replication of the authentic SARS-CoV-2 virus in ACE2-transfected HEK293 cells increased more than four times in the presence of the enhancing antibodies (Fig. 2c). Huh7 cells, with low levels of ACE2^22^, were also susceptible to SARS-CoV-2 infection. SARS-CoV-2 infection of the Huh7 cells was also significantly enhanced by the enhancing antibodies. These data indicated that the enhancing antibodies augment the infectivity of the SARS-CoV-2 virus to ACE2-expressing cells.

### Overlapping epitopes in NTD recognized by the enhancing antibodies

We employed competitive binding assays to identify the epitopes of the infectivity-enhancing antibodies. Surprisingly, all the enhancing antibodies competed for the spike protein with differing efficiencies, suggesting that they recognized similar epitopes on the NTD (Fig. 3a). Next, we generated a series of alanine mutants of NTD with the transmembrane domain (Supplementary Fig. 1a) and analyzed their binding to the enhancing antibodies. Because the 8D2 mAb contained negatively charged amino acids in the heavy chain CDR3, we analyzed the binding of the 8D2 mAb to lysine- or arginine-mutated NTD. The R214A and K187A mutants were not recognized by the 8D2 antibody (Fig. 3b). We then analyzed a series of NTD mutants in which the amino acid residues structurally close to R214 or K187 were mutated to alanine and found that the binding of the enhancing antibodies to the whole spike protein was substantially decreased in W64A, H66A, K187A, V213A, and R214A mutants (Fig. 3b). Similar observations were made with the full-length spike protein with mutations in these residues (Fig. 3c). Moreover, the W64A-H66A and V213A-R214A double mutants reduced the binding of the enhancing antibodies more than the single mutants did. Furthermore, the quadruple mutant of W64, H66, V213, and R214 was not recognized by any enhancing antibody. Significantly, these mutations did not affect the NTD’s recognition by a non-enhancing anti-NTD antibody, 4A8, an anti-RBD antibody, C114, or an anti-S2 antibody, 2454. These data suggested that mutations of these residues in the NTD did not affect the overall conformation of the spike protein (Supplementary Fig. 4).

**Fig. 3.**
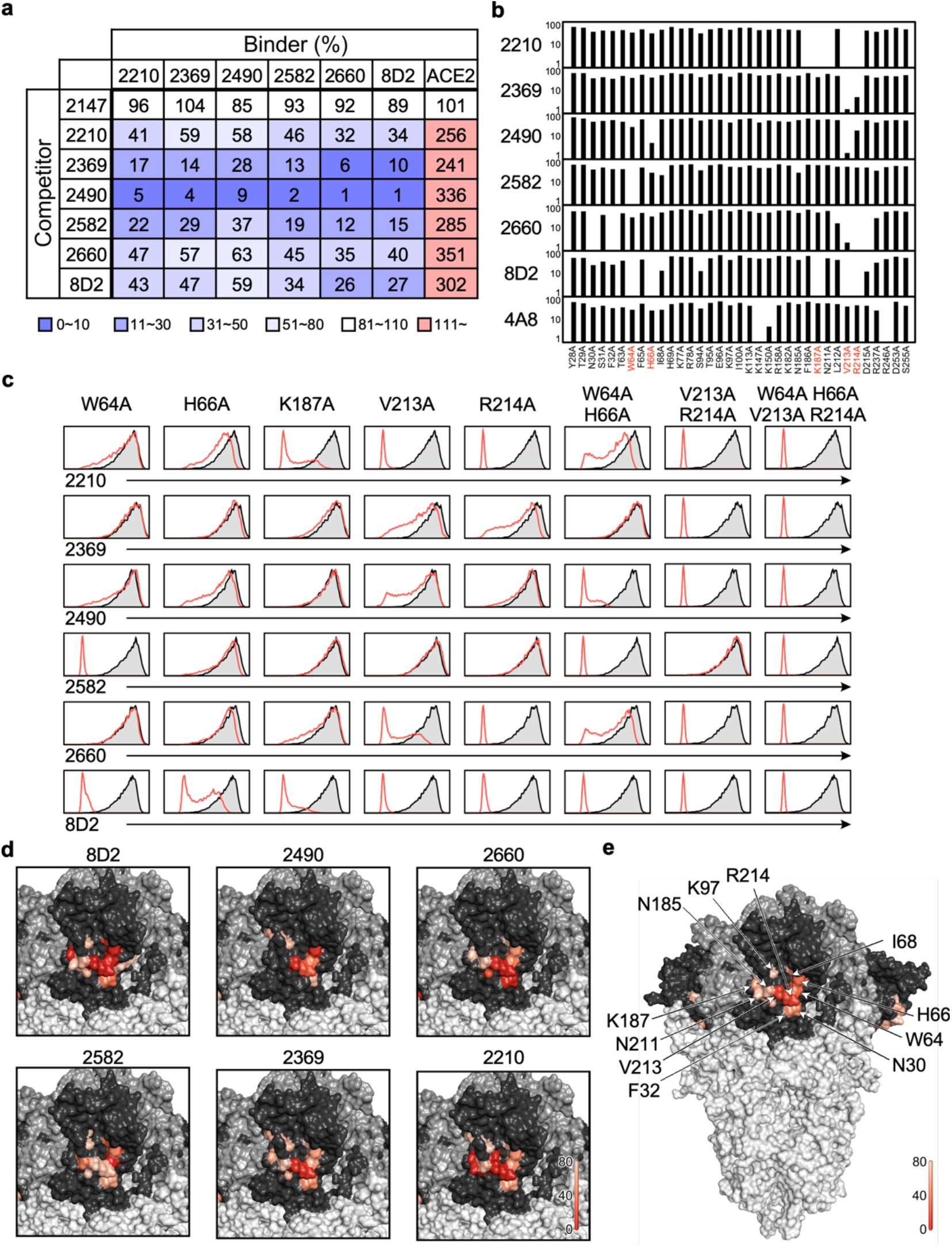
Epitope mapping of the SARS-CoV-2 infectivity-enhancing antibodies. **a**, The binding of the enhancing antibodies (binder) to full-length spike transfectants was analyzed in the presence of the indicated antibodies (competitor). The effect of competitors on ACE2-Fc binding to the spike transfectants was also analyzed. The non-enhancing anti-S2 antibody, 2147, was used as a control. Relative antibody or ACE2 binding lebels observed in the presence of competitor are shown. **b**, Relative antibody binding levels to a series of NTD mutants compared to wild-type NTD are shown. Non-enhancing anti-NTD antibody 4A8 was used as a control. The most affected resideus were shown as red. **c**, The full-length mutant spike proteins were stained with the indicated enhancing antibodies (red line). Staining of wild-type spike were shown as shaded histogram. **d**. Amino acid residues that affected the binding of each enhancing antibodies are shown as a heatmap based on their percent reduction of the MFIs in **b**), with higher reduction indicated by darker shades. NTD: dark grey, RBD: medium grey, other regions: light grey. **e**, The MFIs reduction of the affected residues are averaged across the six antibodies and shown as a heatmap.

Interestingly, the epitopes for the enhancing antibodies were in a narrow area on the NTD (Fig. 3d and 3e). Based on the epitopes for these antibodies, we generated docking models of the antibody–spike protein complex and found that all the enhancing antibodies were similarly bound to NTD (Supplementary Fig. 5). These data suggested that the antibodies binding to these specific epitopes on the NTD may act in a similar manner, possibly facilitating the binding to ACE2 by modulating the local conformation of the NTD.

### Infectivity-enhancing antibodies in hospitalized COVID-19 patients

Because the enhancing antibodies boost SARS-CoV-2 infectivity, we compared the serum levels of the enhancing antibodies in COVID-19 patients to those of uninfected individuals. We utilized a competitive binding assay using recombinant spike protein and fluorescence-labeled 8D2 enhancing antibody to analyze the serum levels of the enhancing antibodies (Fig. 4a). Binding of the 8D2 enhancing antibody to spike protein-coated beads was decreased in the presence of the serum from a COIVD-19 patient but not from an uninfected donor (Fig. 4b).

**Fig. 4.**
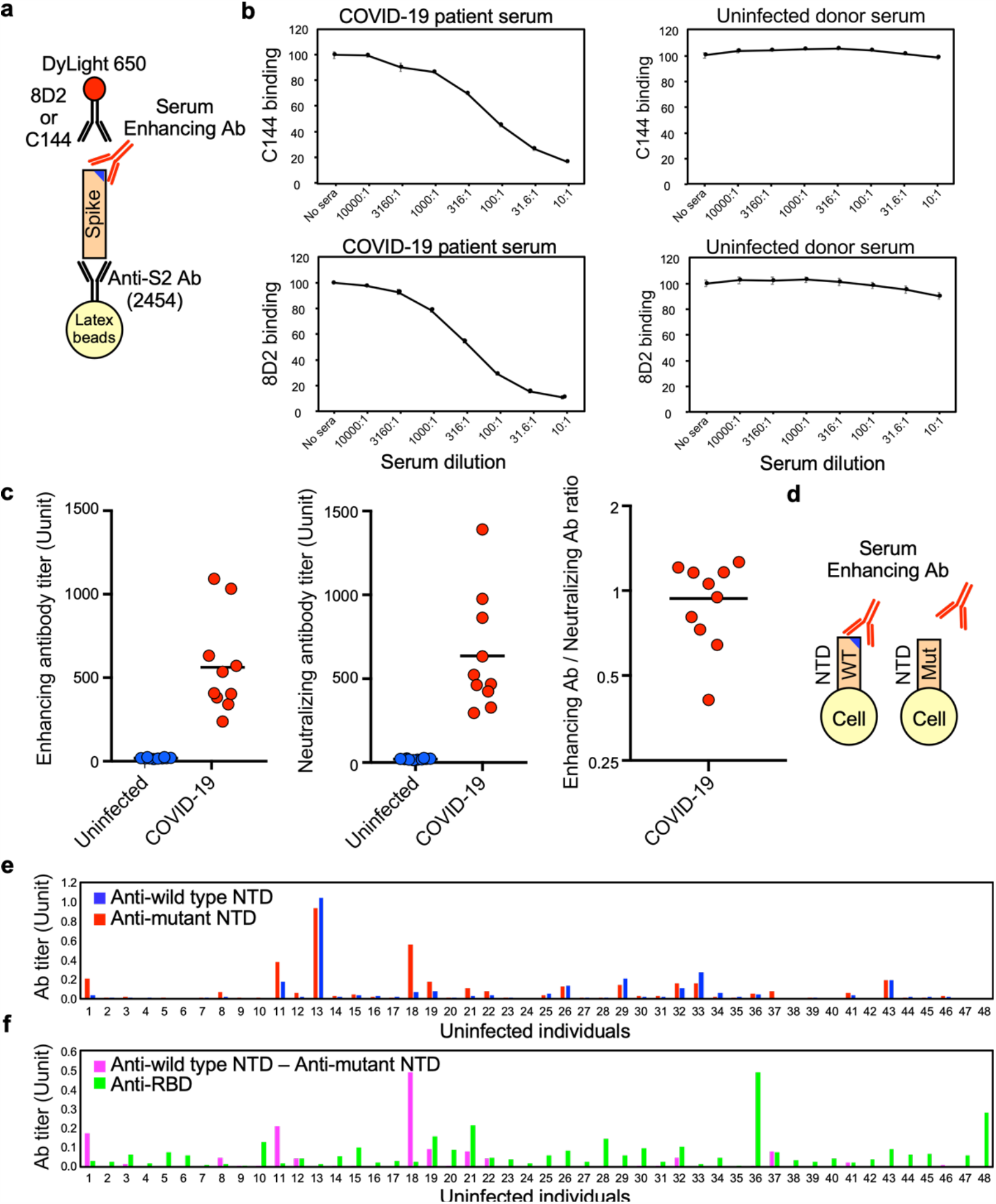
SARS-CoV-2 infectivity-enhancing antibodies in COVID-19 patients and uninfected individuals. **a**, A method of detecting the enhancing or neutralizing antibodies using a competitive binding assay. DyLight 650-labeled 8D2 and C144 were used to detect the enhancing and neutralizing antibodies, respectively. **b**, The binding of the enhancing antibody, 8D2, or the neutralizing antibody, C144, to beads coated with the spike protein was analyzed in the presence of the serially diluted serum of a representative COVID-19 patient or an uninfected donor. **c**. The levels of SARS-CoV-2 infectivity-enhancing antibodies (left) and neutralizing antibodies (middle) in COVID-19 patients (red, n=10) and unifected individuals (blue, n=10) were analyzed using the competitive binding assay. The ratio of enhancing antibodies to neutralizing antibody levels in each COVID-19 patient is shown (right). **d**, The enhancing antibodies were detected by comparing the antibody binding to the wild-type NTD-TM (WT) to the antibody binding to the mutant NTD TM lacking the enhancing antibody epitopes (Mut). **e**, The serum levels of antibodies in uninfected individuals against the wild-type NTD (blue bar) and mutant NTDs whose epitopes for the enhancing antibodies were mutated (red bar). **f**, SARS-CoV-2 infectivity-enhancing antibody titers were calculated by subtracting the antibody levels against the mutant NTD from those against the wild-type NTD in uninfected individuals. Anti-RBD antibody titers (green bar) were analyzed using RBD-TM transfectants (green bar).

We then measured the enhancing antibody titers in COVID-19 patients using this system. The enhancing antibodies and neutralizing antibodies were detected in all the hospitalized COVID-19 patients; however, the ratio of the enhancing antibodies to the neutralizing antibodies was different among them (Fig. 4c). It has been reported that a few uninfected individuals possess antibodies against the SARS-CoV-2 spike protein^23^. Therefore, we investigated whether the uninfected individuals carried the enhancing antibodies. Because the competitive binding assay used to analyze COVID-19 patients was not sensitive enough to detect low levels of the enhancing antibodies in uninfected individuals, we measured the levels of the enhancing antibodies by comparing a serum antibody binding to wild-type SARS-CoV-2 NTD and to mutant SARS-CoV-2 NTD (W64A, H66A, K187A, V213A, and R214A) that was not bound by enhancing antibodies (Fig. 4d). Some donors, such as #13, possessed antibodies bound to both the wild-type and mutant NTD, suggesting that they lacked the enhancing antibodies (Fig. 4e). By contrast, other donors, such as #18, produced antibodies specific to wild-type NTD, suggesting that they had the enhancing antibodies. Some donors also possessed low levels of anti-RBD antibodies, but there was no correlation between the titers of the anti-RBD antibody and the enhancing antibody (Fig. 4f). These data indicate that antibody responses against specific sites of the SARS-CoV-2 spike protein vary significantly between individuals.

## Discussion

Antibody-dependent enhancement (ADE) of viral infection has been reported for some viruses such as dengue virus^24^, feline infectious peritonitis (FIP) coronavirus^25,26^, SARS^27,28^, and MERS^24^. Binding of the Fc receptor to anti-virus antibodies complexed with virions has been thought to be involved in ADE^13^. However, Fc receptor-mediated ADE is restricted to the infection of Fc receptor-expressing cells. In this study, we found a non-canonical, Fc receptor-independent ADE mechanism. The antibodies against a specific site on the NTD of the SARS-CoV-2 spike protein were found to directly augment the binding of ACE2 to the spike protein, consequently increasing SARS-CoV-2 infectivity. Because the Fc receptor is not involved in this new type of ADE, the facilitation of infection by the enhancing antibodies could be involved in the SARS-CoV-2 infection of a broad range of cells.

The spike protein RBD is quite flexible, and ACE2 preferentially binds to the open form of the RBD^29,30^. Most structural studies of the wild-type spike protein exhibited one RBD in the open conformation. By contrast, two or three RBDs were observed in the open formation in the D614G mutant, suggesting that the RBD conformation plays a vital role in the infectivity of SARS-CoV-2^15,29^. The RBD is structurally associated with the NTD of a neighboring chain^6,31^. Since the conformations of viral proteins can be altered significantly upon antibody binding^32,33^, the binding of enhancing antibodies to a specific site on the NTD may modulate the NTD-RBD interchain interaction, thus increasing the open conformation of RBD.

All the enhancing monoclonal antibodies used in this study were derived from COVID-19 patients. Notably, the enhancing antibodies were detected at similar levels to those of the neutralizing antibodies in the sera of hospitalized COVID-19 patients. In addition, the ratio of the two types of antibodies differed among COVID-19 patients. Therefore, the extent of production of enhancing antibodies relative to that of anti-RBD neutralizing antibodies could be a factor involved in COVID-19 disease progression.

Surprisingly, a few uninfected individuals possess antibodies that recognize the infectivity-enhancing site of the NTD. Although anti-spike antibodies are often detected in uninfected individuals, most of them are directed against the S2 subunit^23^. The sequence homology of the NTD to common human coronaviruses is low compared to the S2 subunit. In particular, the antibody epitopes on the enhancing site are not conserved among other coronaviruses. Although it is unclear how the enhancing antibodies were produced in some uninfected individuals, the production of the enhancing antibodies may be boosted by SARS-CoV-2 infection or vaccination. Therefore, spike proteins that lack the antibody epitopes at the enhancing site might have to be considered to vaccinate individuals with pre-existing enhancing antibodies. Transfusion of plasma from recovered COVID-19 patients has been proposed as a possible treatment for COVID-19^34^. Because the ratio of neutralizing antibodies to enhancing antibodies in serum differs among COVID-19 patients, the plasma level of the enhancing antibodies in the donor may affect the treatment’s effectiveness. Therefore, the function of the enhancing antibodies in SARS-CoV-2 infection has to be further analyzed in both COVID-19 patients and healthy individuals.

## Methods

### Human Samples

The collection and use of human sera and synovial tissues were approved by Osaka University (2020-10 and 19546), Kobe City Medical Center General Hospital (200924), and Osaka South Hospital (2-28). Written informed consent was obtained from the participants according to the relevant guidelines of the institutional review board. The diagnoses of SARS-CoV-2 were PCR-based. The sera from uninfected humans were taken before June 2019 (George King Bio-Medical). All the sera of the SARS-CoV-2 patients were treated with 2% CHAPS for 30 min at room temperature to inactivate the remaining virus.

### Cell lines and cell culture

HEK293T cells (RIKEN Cell Bank), Huh7 cells (the National Institute of Infectious Diseases) and TMPRSS2-expressing VeroE6 cells (Japanese Collection of Research Bioresources Cell Bank, JCRB1819) were cultured in DMEM (GIBCO, 11995-073) supplemented with 10% FBS (Biological Industries, USA), penicillin (100 U/mL), and streptomycin (100 μg/mL) (Nacalai, Japan) and cultured at 37°C in 5% CO2. The Expi293 cells (Thermo) were cultured with the Expi293 medium. The cells were routinely checked for mycoplasma contamination.

### Genes and plasmids

The SARS-CoV-2 spike gene (NC_045512.2) was prepared by gene synthesis (IDT). The sequences encoding the spike protein lacking C-terminal 19 amino acids (amino acids 1–1254) were cloned into pME18S expression vector. NTD (amino acids 14–333), RBD (amino acids 335–587), and S2 (amino acids 588–1219) were separately cloned into a pME18S expression vector containing a SLAM signal sequence and a PILRα transmembrane domain^35^. A series of alanine mutants were introduced into the SARS-CoV-2 spike protein using the QuickChange mutagenesis kit or the QuickChange multi-mutagenesis kit (Agilent). The primers for mutagenesis were designed on Agilent’s website (https://www.agilent.com/store/primerDesignProgram.jsp). The cDNA encoding ACE2 (NM_021804.3) was cloned into a pMxs retrovirus vector. The mouse ACE2-IgG2a Fc fusion protein was prepared by cloning the sequence encoding the extracellular domain of *Ace2* (amino acid residues 20–740) into the pCAGGS expression vector containing the SLAM signal sequence and the sequence encoding the mouse IgG2a Fc domain as previously described^36^. The sequence encoding the spike protein’s extracellular domain with a foldon and His-tag at the C-terminus^5^ was cloned into a pcDNA3.4 expression vector containing the SLAM signal sequence. Also, mutations D614G, R686G R687S R689G, K986P, and V987P were introduced using a Quick change multi-mutagenesis kit (Agilent) for spike protein stabilization^15^. The DNA sequences of these constructs were confirmed by sequencing (ABI3130xl).

### Transfection

A pME18S expression plasmid containing the full-length or subdomain spike protein was transiently transfected into HEK293T cells using PEI max (PolyScience); the pMx-GFP expression plasmids were used as the marker of transfected cells. ACE2 was stably transfected into HEK293T cells using the pMxs retrovirus vector. Briefly, pMxs-ACE2 and amphotropic envelope vectors were co-transfected into PLAT-E packaging cells. The cell culture supernatants containing the retroviruses were harvested 48 hours later and premixed with DOTAP (Roche, Switzerland) before spin transfection at 2400 rpm and 32°C for 2 hours. Afterward, the ACE2-expressing HEK293T cells were purified using a cell sorter (SH800S, Sony).

### Anti-spike antibodies from COVID-19 patients

The V regions of anti-SARS-CoV-2 spike mAb from COVID-19 patients were synthesized according to the published sequence (IDT)^9-11,14^. The cDNA sequences of the variable regions of the heavy chain and light chain were cloned into a pCAGGS vector containing sequences that encode the human IgG1 or kappa constant region. The IgG with the murine IgG2a constant region was used for the competitive binding assay. The pCAGGS vectors containing sequences encoding the immunoglobulin heavy chain and light chain were co-transfected into HEK293T cells or Expi293 (Thermo) cells, and the cell culture supernatants were collected according to the manufacturer’s protocols. Recombinant IgG was purified from the culture supernatants using protein A Sepharose (GE healthcare) except for Fig. 1a. The concentration of unpurified IgG in the cell culture supernatants used in Fig. 1a was measured using the protein A-coupled latex beads (Thermo A37304) and APC-labeled anti-human IgG F(ab’)2 antibodies (Jackson) against IgG standards of known concentration. The concentration of purified IgG was measured at OD280. DyLight 650 (Thermo)-labeled 8D2 or C144 mAbs was used to detect the enhancing antibodies or neutralizing antibodies, respectively. Protein A-purified 8D2^14^ and C144^10^ antibodies were labeled using DyLight 650 amine-reactive dye according to the manufacturer’s protocol.

### Antibodies and recombinant proteins

Mouse anti-human ACE2 mAb (R&D Systems, USA), rat anti-Flag mAb (L5, Biolegend), allophycocyanin (APC)-conjugated donkey anti-mouse IgG Fc fragment Ab, APC-conjugated anti-human IgG Fc fragment specific Ab, APC-conjugated anti-rat IgG specific Ab, and APC-conjugated streptavidin (Jackson ImmunoResearch, USA) were used. The pCAGGS vector containing a sequence that encodes the ACE2-mouse IgG2a Fc fusion protein was transfected into the HEK293T cells. Recombinant ACE2-Fc fusion protein was purified from the cell culture supernatants with the protein A Sepharose (GE Healthcare). The purified ACE2-mouse IgG2a Fc fusion protein was biotinylated with Sulfo-NHS-LC-Biotin (Thermo), followed by buffer exchange with a Zeba spin desalting column. The pcDNA3.4 expression vector containing the sequence that encodes the His-tagged extracellular domain of the spike protein was transfected into Expi293 cells; then, the His-tagged spike protein produced in the culture supernatants was purified with a Talon resin (Clontech).

### The flow cytometric analysis of antibodies

The plasmid expressing the full-length SARS-CoV-2 spike protein, Flag-NTD-PILR-TM, Flag-RBD-PILR-TM, Flag-S2-PILR-TM, or mutated spike proteins were co-transfected with the GFP vector into HEK293T cells. The transfectants were incubated with the mAbs, followed by APC-conjugated anti-human IgG Ab. Then, the antibodies bound to the stained cells were analyzed using flow cytometers (Attune™, Thermo; FACSCalibur BD bioscience). Antibody binding to the GFP-positive cells were shown in the figures using a FlowJo software (BD bioscience).

### ACE2 binding assay

Because SARS-CoV-2 spike protein contains intracellular retention signal at C-terminus^37,38^, we used spike protein lacking C-terminal 19 amino acids for ACE2 binding assay. C-terminal deleted spike protein and GFP were cotransfected into HEK293T cells and the transfectants were mixed with various concentrations of anti-spike antibodies at 4°C for 30 min, followed by incubation with a biotinylated-ACE2-Fc fusion protein at 1 μg / ml and 4°C for 30 min. Thereafter, the ACE2 bound to the spike protein was detected using APC-conjugated streptavidin (2.5 μg/mL). The amount of ACE2 on GFP-positive cells was analyzed by a flow cytometer.

### SARS-CoV-2 spike-pseudotyped virus infection assay

The HEK293T cells were transiently transfected with expression plasmids for the SARS-CoV-2 spike protein lacking C-terminal 19 amino acids^37,38^. Twenty-four hours post transfection, VSV-G-deficient VSV carrying a GFP gene complemented in trans with the VSV-G protein was added for a two-hour incubation. The cells were then carefully washed with DMEM media without FBS and incubated with DMEM with FBS at 37°C in 5% CO_2_ for 48 hours. The supernatant containing the pseudotyped SARS-CoV-2 virions was harvested and aliquoted before storage at −80°C. The virus titers were determined using TMPRSS2-expressing VeroE6 cells. The pseudotyped SARS-CoV-2 virus was pre-incubated with anti-NTD monoclonal antibodies for 30 minutes and mixed with the ACE2-expressing cells. Twenty-four hours later, dead cells were stained with propidium iodide (Sigma), and the proportions of GFP-expressing cells in the living cells were analyzed by flow cytometry.

### The Authentic SARS-CoV-2 infection assay

The authentic SCoV-2 infection assay was carried out in a Biosafety Level 3 laboratory. The virus strain was obtained from the Kanagawa Prefectural Institute of Public Health (KNG19-020). The stock virus was amplified in *TMPRSS2*-expressing VeroE6 cells. The virus stock (6.25 log TCID50) was prediluted 1: 100 in DMEM supplemented with 2% FBS before being incubated with the cells at 1 × 10^5^ cells/mL at 37°C in 5% CO_2_ for 3 hours. The virus was then removed by washing the cells with DMEM; the cell culture supernatant was collected for control. The cells were incubated for another 2 days. The cell culture supernatant was collected for viral RNA extraction using the QIAamp viral RNA extraction kit (Qiagen, Germany) and subsequently quantified using real-time PCR using the N2 primer set (AGCCTCTTCTCGTTCCTCATCAC and CCGCCATTGCCAGCCATTC).

### The competitive binding assay

The binding of the enhancing antibodies containing a mouse IgG2a constant region (1 μg/ml) to full-length spike transfectants was analyzed in the presence of the human antibodies (10 μg/ml). The effect of competitors on ACE2-Fc binding to the spike transfectants was also analyzed. The non-enhancing anti-S2 antibody, 2147, was used as a control. Relative mean fluorescence intensities observed in the presence of competitor antibodies were calcularated.

### The docking model of the enhancing antibodies and spike protein

A homology model of the SARS-CoV-2 prefusion spike trimer was built using a template-based method. The prefusion spike trimer structure (PDB ID: 7jji) in the all-RBD-down status was used as the main structural template. To model missing residues within the S2 domain, fragments were sampled from other SARS-CoV-2 spike trimer structures. The best fragment template was selected as the one with the lowest RMSD within flanking regions of the main template and the complete structure was constructed using MODELLER^39^. We investigated possible binding modes of the enhancing antibodies through antibody modelling followed by docking. Antibody structures were modelled by Repertoire Builder^40^ and docked onto the NTD using the HADDOCK2.4 webserver^41^ using observed epitope residues as constraints. The top-scoring docked poses were rendered on the full-length spike model using PyMOL. Residues identified by alanine scanning to affect enhancement were represented as a heatmap onto the surface of the spike model.

### The analysis of the enhancing and neutralizing Ab titers

The 4 µm aldehyde/sulfate latex beads (Thermo A37304) were coated with protein A-purified anti-S2 mAb (2454)^11^ and then incubated with recombinant spike protein at 7 μg/ml to capture the recombinant SARS-CoV-2 spike protein on the latex beads. The latex beads containing recombinant SARS-CoV-2 spike were then incubated with 1:30 diluted serum from COVID-19 patients or uninfected individuals and stained with DyLight 650-labeled 8D2 mAb at 3 μg/ml) or C144 mAb at 3 μg/ml. After fixation with 4% paraformaldehyde (Nacalai Tesque), the beads were analyzed using flow cytometry. We defined the high levels of the enhancing and neutralizing antibody titers in a COVID-19 patient’s serum as 1,000 units and used the serum as a standard to calculate antibody titers of other serum samples.

### Detection of enhancing antibody in uninfected individuals

The plasmids expressing the wild-type NTD-TM, which was recognized by the enhancing antibodies, and the NTD-TM mutants (W64A, H66A, K187A,V213A, and R214A) not recognized by all enhancing antibodieswere transfected into HEK293T cells with the GFP vector as a transfection marker. The transfectants were mixed with 1:100 diluted serum from uninfected individuals, and the bound antibodies were detected with the APC-labeled anti-human IgG Fc antibody. The 4A8 anti-NTD antibody^14^ was used as a standard to calculate the relative concentration of the antibody against NTD. The stained cells were analyzed by a flow cytometer. The level of serum antibodies specific to the wild-type or mutant NTD was calculated by subtracting the mean fluorescence intensity of the antibodies bound to the GFP-negative cells from those of the GFP-positive cells. The level of the enhancing antibodies was calculated by subtracting antibody levels against the mutant NTD from those against the wild-type NTD. Similarly, the relative level of the anti-RBD antibody was measured using RBD-TM transfectants and C144 as a standard. One μg/ml of 4A8 and C144 was defined as 1 unit.

### Data and statistical analysis

FlowJo version 10.7 (BD Biosciences, USA) was used to analyze the flow cytometry data, and Graphpad Prism version 7.0e was used for graph generation and statistical analysis.

## Acknowledgements

Soh W.T. is supported under the Kishimoto Foundation Fellowship. We would like to thank Hiroshi Honda for technical assistance and Chikako Kita for administrative assistance. This work was supported by JSPS KAAKENHI under Grant Numbers JP18H05279 and JP19H03478, MEXT KAKENHI under Grant Number JP19H04808, Japan Agency for Medical Research and Development (AMED) under Grant Numbers 19fk0108161h0001, 20nf0101623h0201, and 20nk0101602h0201.

## Data availability

Source data for figures are provided with the paper.

## Author contributions

Y.L., Y.S., M.O., Y.M., D.M.S., S.T., and H.A. conceived experiments. Y.L., W.T.S., A.T, A.A., S.M., E.E M.Y., S.T., N.E.E., C.O., S.T., K.K., H.J. W.N., M.K., and H.A. performed experiments. A.N., N.A., N.H., S.H., K.T, K.O. collected patients’ sera. Y.L. and H.A. wrote the manuscript. All authors read, edit, and approved the manuscript.

---

**Supplementary Fig. 1.**
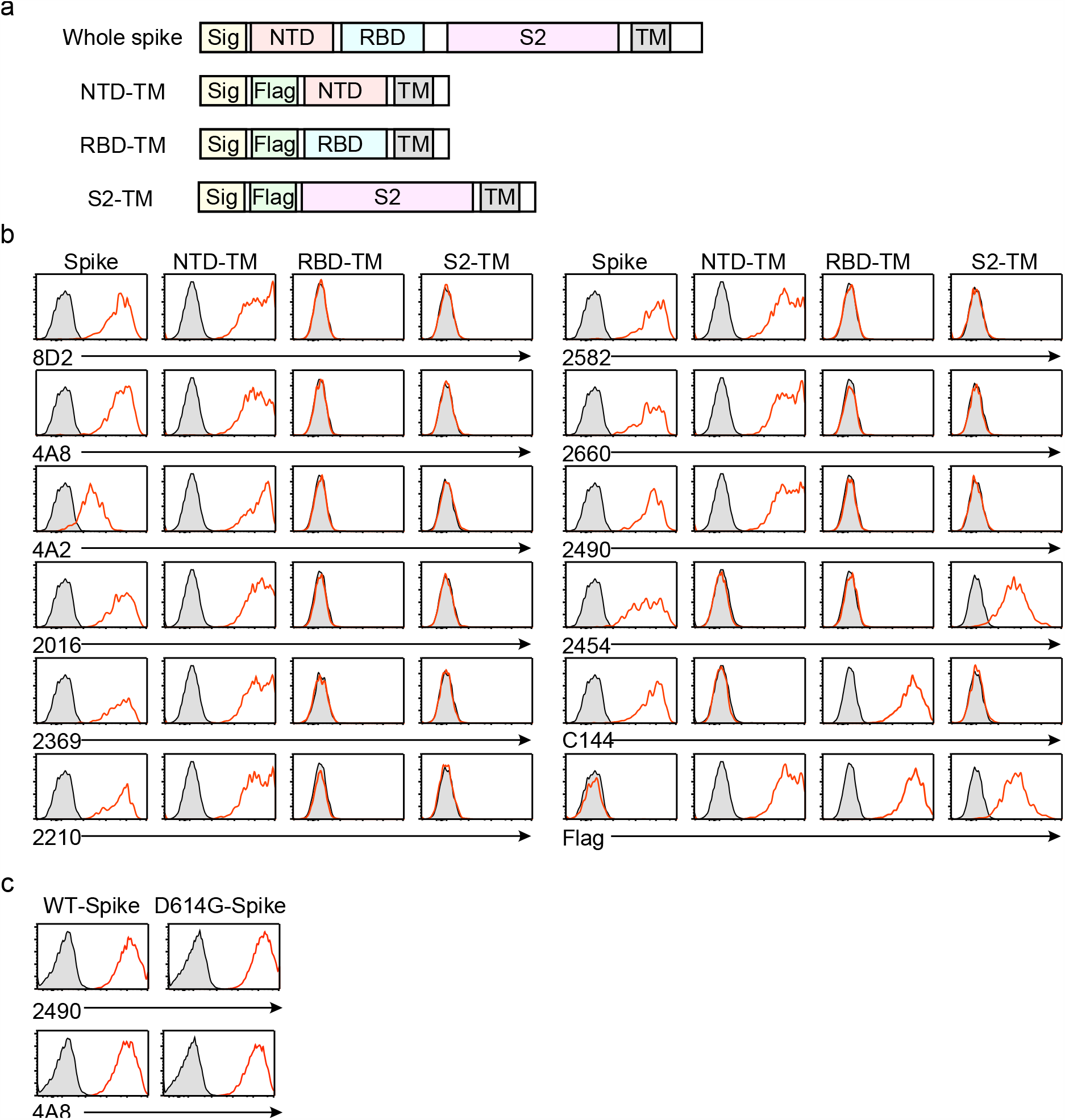
The specificities of the antibodies used in this study. **a**, The expression constructs used to analyze the specificity of the anti-spike antibodies. **b**, The plasmids encoding the full-length spike, Flag-NTD-TM, Flag-RBD-TM, and Flag-S2-TM were cotransfected separately with a GFP vector into HEK293 cells. The transfectants were then stained with the indicated antibodies. The antibodies bound to the transfectants were detected with APC-labeled secondary antibodies. The fluoresce intensities of APC on the GFP-expressing cells are shown (red line). Control stainings were shown as shaded histogram. **c**, The plasmids expressing the wild-type spike protein and the D614G mutant were cotransfected separately with a GFP vector into HEK293T cells. The transfectants were stained with the anti-NTD infectivity-enhancing antibody 2490 and anti-NTD non-enhancing antibody 4A8. The fluorescence intensities of APC on the GFP-expressing cells are shown (red line). Control stainings were shown as shaded histogram.

**Supplementary Fig. 2.**
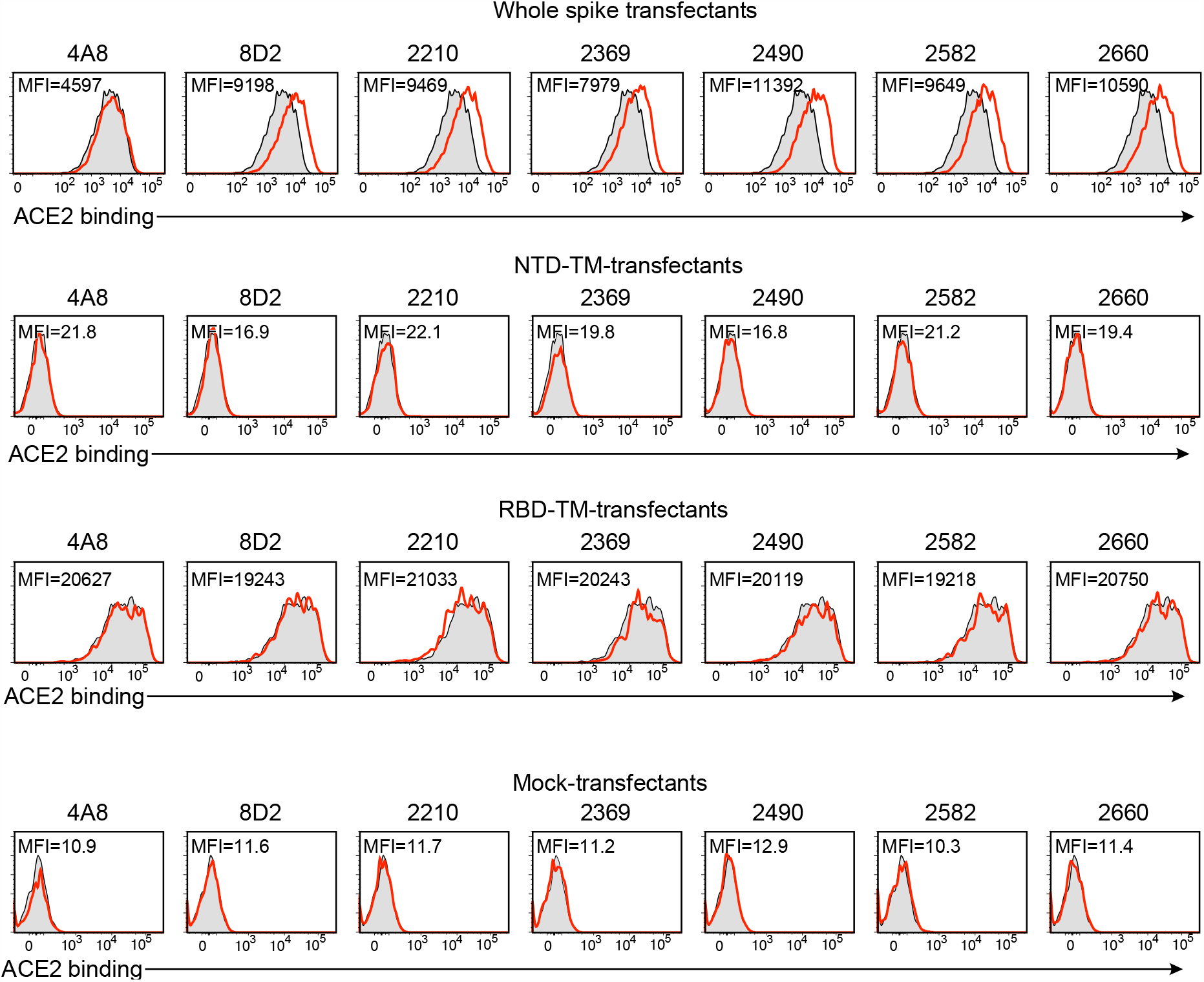
The enhanced binding of ACE2 to the spike protein by specific anti-NTD antibodies. The plasmids expressing the full-length spike, NTD-TM, RBD-TM, and mock were transfected into HEK293T cells with the GFP vector, and the transfectants were mixed with the indicated anti-NTD antibodies at 10 μg/ml. 4A8 is a non-enhancing antibody and the remaining is enhancing antibodies. Transfectants not mixed with antibodies were used as a control (shaded histogram). Afterward, the cells were stained with biotin-labeled ACE2-Fc fusion protein, followed by APC-labeled streptavidin. The fluorescence intensities of APC on the GFP-expressing cells are shown (red line). Mean fluorescent intensities (MFI) of red lines were shown in the figure.

**Supplementary Fig. 3.**
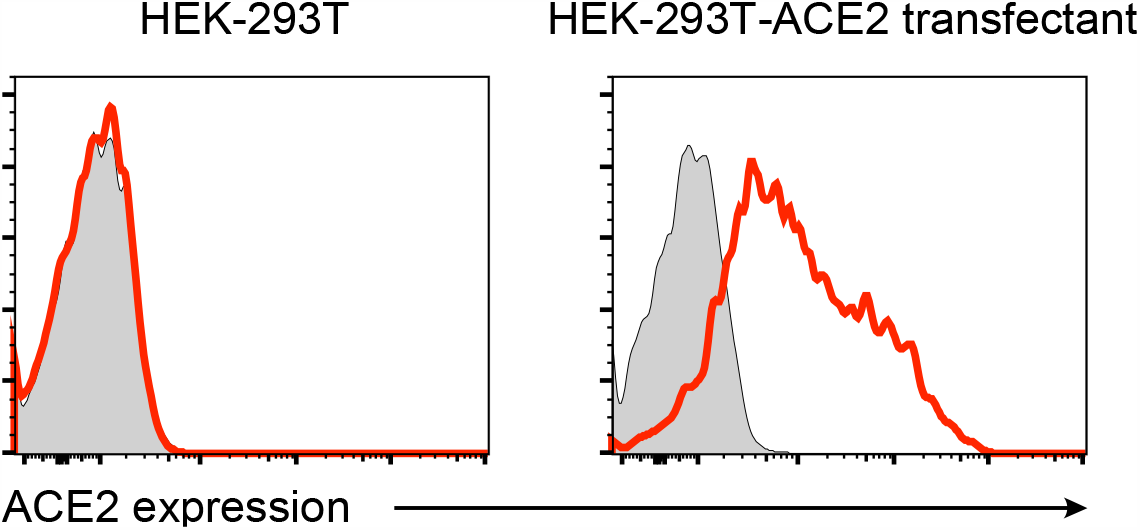
Cell surface levels of ACE2 on HEK293-ACE2 stable transfectants. Parental HEK293T cells and ACE2-transfected HEK293T cells were stained with anti-ACE2 mAb (red line). Control stainings were shown as shaded histogram.

**Supplementary Fig. 4.**
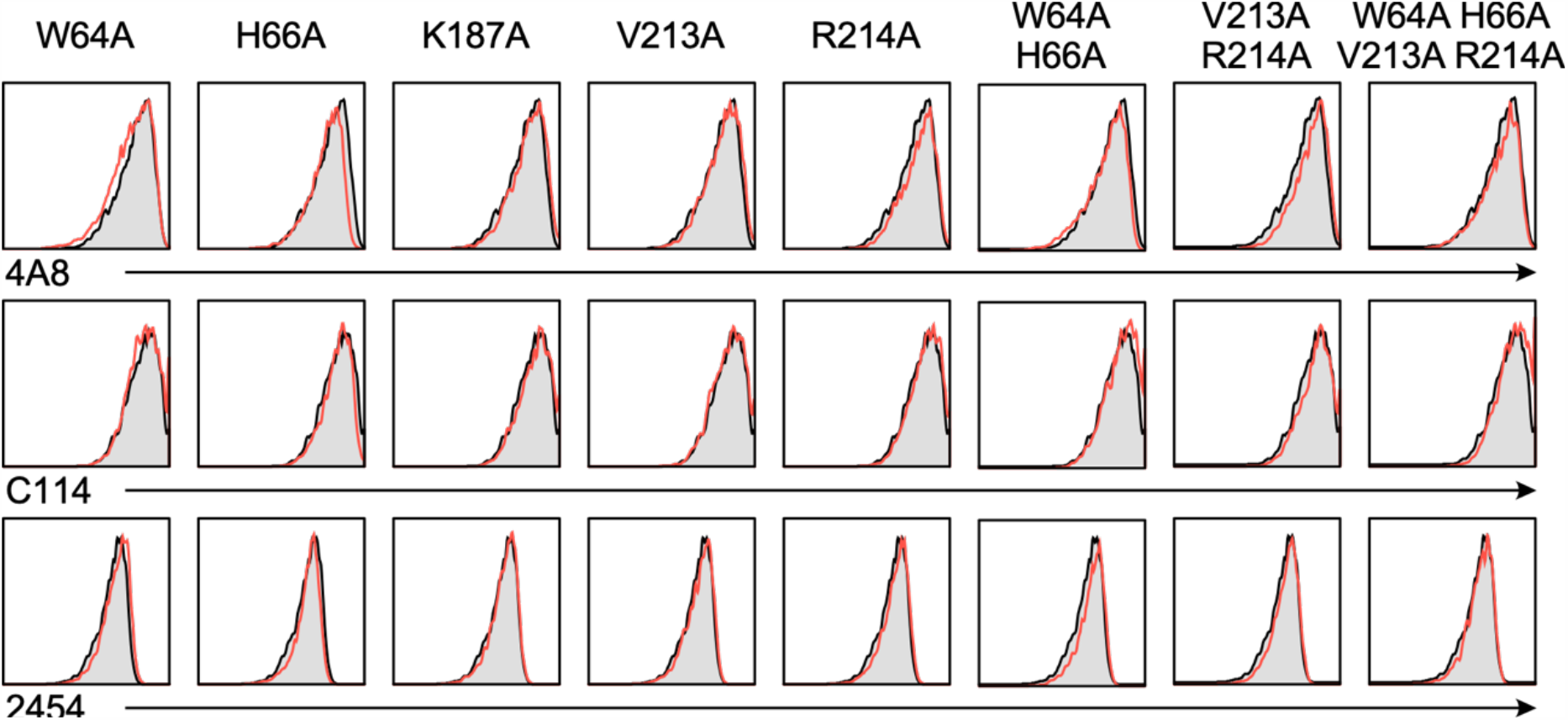
The binding of anti-spike antibodies against spike mutants with mutated antibody epitopes in the SARS-CoV-2 infectivity site. The plasmids expressing the full-length spike proteins with alanine mutations at the indicated amino acid residues were transfected separately with the GFP vector into HEK293 cells, and the binding of 4A8 (anti-NTD non-enhancing antibody), C144 (anti-RBD antibody), and 2454 (anti-S2 antibody) against the GFP-positive cells were analyzed (red line). Control stainings were shown as shaded histogram.

**Supplementary Fig. 5.**
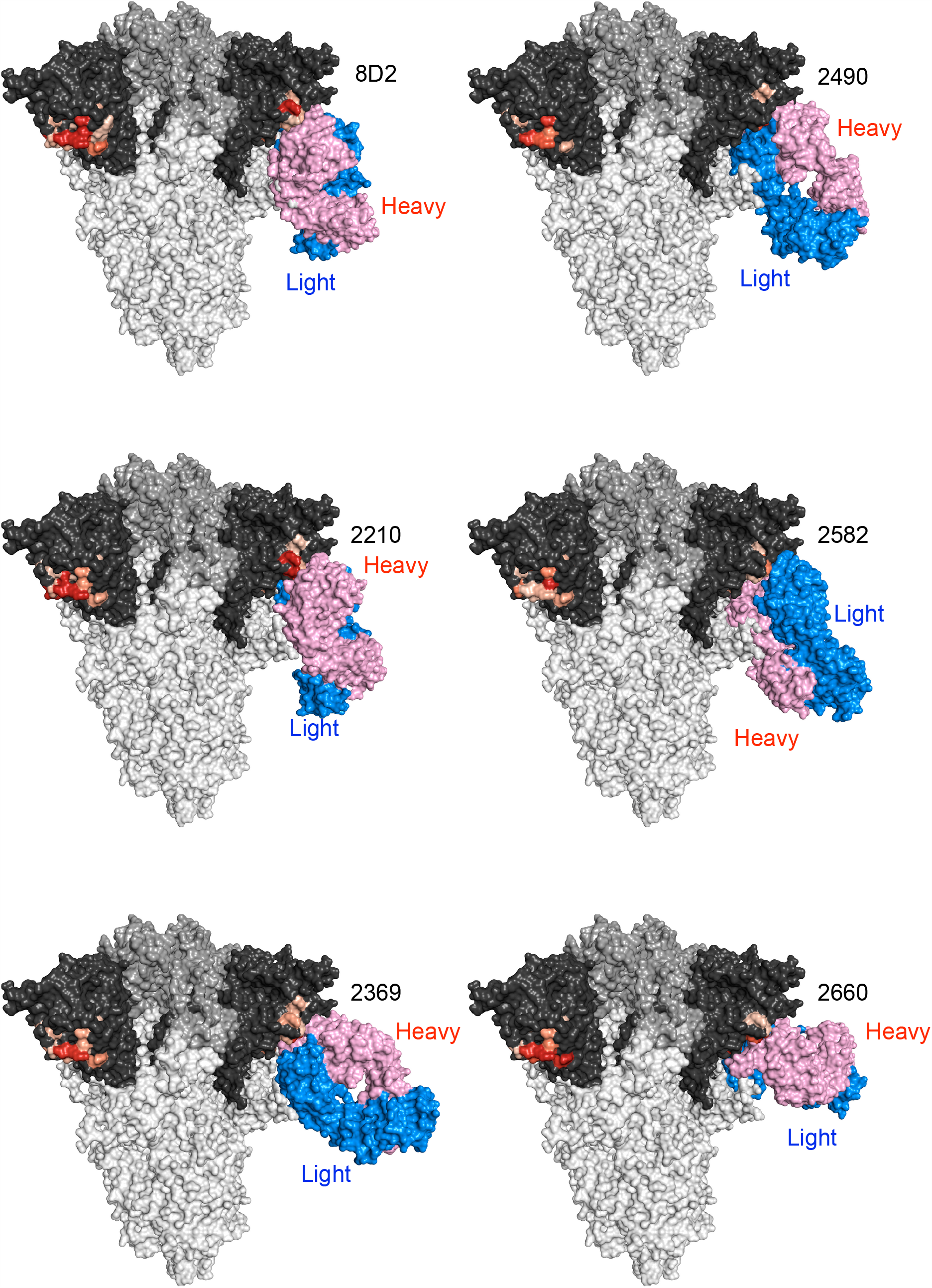
Docking model of SARS-CoV2 infectivity enhancing antibodies with spike protein. Each SARS-CoV2 infectivity enhancing antibody was docked onto the spike protein as described in Methods.

